# 3D-Printing and upper-limb prosthetic sockets; promises and pitfalls

**DOI:** 10.1101/2020.09.21.306050

**Authors:** Jennifer Olsen, Sarah Day, Sigrid Dupan, Kianoush Nazarpour, Matthew Dyson

## Abstract

Modernising the way upper-limb prosthetic sockets are made has seen limited progress. The casting techniques that are employed in clinics today resemble those developed over 50 years ago and there is still a heavy reliance on manual labour. Modern manufacturing methods such as 3D scanning and printing are often presented as ready-to-use solutions for producing low-cost functional devices, with public perceptions being largely shaped by the superficial media representation and advertising. The promise is that modern socket manufacturing methods can improve patient satisfaction, decrease manufacturing times and reduce the workload in the clinic. However, the perception in the clinical community is that total conversion to digital methods in a clinical environment is not straightforward. Anecdotally, there is currently a disconnect between those developing technology to produce prosthetic devices and the actual needs of clinicians and people with limb difference. In this paper, we demonstrate strengths and drawbacks of a fully digitised, low-cost trans-radial diagnostic socket making process, informed by clinical expertise. We present volunteer feedback on the digitally created sockets and provide expert commentary on the use of digital tools in upper-limb socket manufacturing. We show that it is possible to utilise 3D scanning and printing, but only if the process is informed by expert knowledge. We bring examples to demonstrate how and why the process may go wrong. Finally, we provide discussion on why progress in modernising the manufacturing of upper-limb sockets has been slow yet it is still too early to rule out digital methods.

## 1 Introduction

Media reports have contributed to misconceptions about the role of digital technology such as 3D printing in the manufacture of upper-limb prosthetics [13], and propagated the perception that modern technology can replace traditional techniques in clinics. It is known within the clinical community that this is far from reality. Limited elements of digital manufacturing were introduced to clinics over thirty years ago [33]. Some clinics have adopted a semi-digital workflow [43], yet a reliable fully digitised method is yet to materialise in upper-limb socket manufacturing. This paper aims to:

- Investigate whether a fully digital *hands-off* trans-radial socket making process is possible, using low-cost optical scanning and 3D printing.
- Present volunteer feedback on digitally created sockets.
- Provide expert commentary on the use of digital tools in upper-limb socket manufacturing.
- Discuss why modernisation of upper-limb socket manufacturing has been slow and the contributions digital methods could still make.

Traditional methods of making sockets, which comprise primarily casting, modification and lamination, can produce comfortable sockets within a reasonable time frame [11]. However, patient feedback has identified lack of comfort as a consistent cause of dissatisfaction and prosthesis abandonment [4, 21, 28, 38]. Amputees must make several visits to specialist clinics, due to the nature of plaster casting [11, 27, 34], and can wait around 2-5 weeks to receive their socket [5, 19, 32, 43]. In the US it is estimated that amputees make an average of nine visits to the clinic per year [31]. Conventionally, no physical or digital record of the cast is kept [11, 25]. Small, manual modifications have a significant impact on the final fit of the socket. Traditional socket making is destructive, limb captures are not retained throughout the process [11] and some steps have no recourse for rectifying mistakes. In the event of irreversable errors the entire manufacturing process must be repeated; requiring the patient to attend the clinic again [11, 34], with accumulating inconvenience for all involved. As such, from the clinical point of view, replacing a socket entails significant labour, is time-consuming [8, 27] and has limited financial return.

The last decade has seen an explosion of interest in digital manufacturing methods [41, 45]. 3D scanning and printing have been introduced as potential technologies that can radically enhance the field of prosthetics [7, 23, 39, 41, 45]. 3D scanners, including low-cost options such as smartphone scanning, can accurately capture the volume and geometry of residual limbs and existing sockets [12, 37] and can eliminate the need for plaster casting and mould disposal [15]. The original and any follow-up scans or digital modifications can be preserved, providing a permanent record of a patient’s limb [11, 25]. In addition, it allows the use of a variety of materials for socket manufacturing, including antibacterial skin-safe filaments [48]. Despite such potential, 3D printing has been used primarily for grasping devices [20]. Whilst 3D printed sockets for lower limb have shown early success, in terms of comfort and reduction of contact pressure [7], experts have argued that 3D printing is unlikely to have a significant influence on upper-limb socket manufacturing [45].

## 2 Method

### 2.1 Ethics

The local ethics committee at Newcastle University approved this study (Ref: #16602/2018).

### 2.2 Participants

Six participants with trans-radial limb absence took part in this experiment. Five were acquired amputees and one had congenital limb deficiency. Three of the six participants reported pain and phantom limb sensations to varying degrees of severity. Further participants details can be found in Table 1.

**Table 1:** Table shows participant gender (self-identified); age-range; cause (A: Amputation, C: Congenital); prosthesis type (P: Passive, BP: Body-Powered, M: Myoelectric) and frequency of use; notes (PLS: Phantom Limb Sensations, P: Pain).

| Participant | Gender | Age | Cause | Prosthesis type | Prosthesis Use | Notes |
| --- | --- | --- | --- | --- | --- | --- |
| P1 | M | 50-60 | A | P/BP | Occasional | PLS & P |
| P2 | F | 40-50 | A | P/BP | Occasional | PLS (mild) & P |
| P3 | M | 30-40 | A | M | Daily | - |
| P4 | M | 20-30 | C | P | Occasional | - |
| P5 | M | 60+ | A | P/BP | Frequent | PLS (constant) & P |
| P6 | M | 60+ | A | P/BP | Occasional | - |

### 2.3 Traditional build of sockets

In order to digitally replicate the traditional method of manufacturing sockets, an upper-limb socket making method taught at an ISPO certified P&O training school was observed. Despite processes varying between clinics, the general steps are as follows:

1. **Patient History:** Initial consultation with the patient to gather background information and identify the sensitive and painful areas of the limb.
2. **Limb preparation:** Marking areas of interest on the limb using an indelible pencil, either directly to the skin or over a thin sock, e.g. the olecranon, epicondlyes, and any sensitive areas. Fig. 1 shows a diagram displaying key terminology.
3. **Limb shape capture:** Wrapping the limb in plaster soaked bandages, whilst the prosthetist moulds the plaster and applies pressure over areas that will assist with suspension and stability. The cast is allowed to dry and then removed to be filled with plaster to create a positive model of the limb.
4. **Initial modification:** The positive model is adjusted and smoothed by the prosthetist to ensure a correct socket fit and pressure distribution.
5. **Diagnostic socket manufacturing:** A transparent plastic sheet is vacuum formed around the model to create a *diagnostic socket*, aka test or check socket.
6. **Further modification:** The diagnostic socket is fitted to the wearer and checked for fit, comfort and suspension. Adjustments may be made using a heat gun or notes taken for further positive model modification.
7. **Modified positive model creation:** The socket is filled with plaster to create a final positive model of the limb.
8. **Socket fabrication:** The final socket is made, generally using lamination, where several layers of cotton, nyglass and other soft textiles are set with resin around the cast, or vacuum forming using a thermoplastic.
9. **Final additions:** The final socket is fitted to the patient and checked. A second lamination is applied on top of the first, which forms the outer layer of the prosthesis and adds functional details e.g. the wrist.

**Figure 1:**
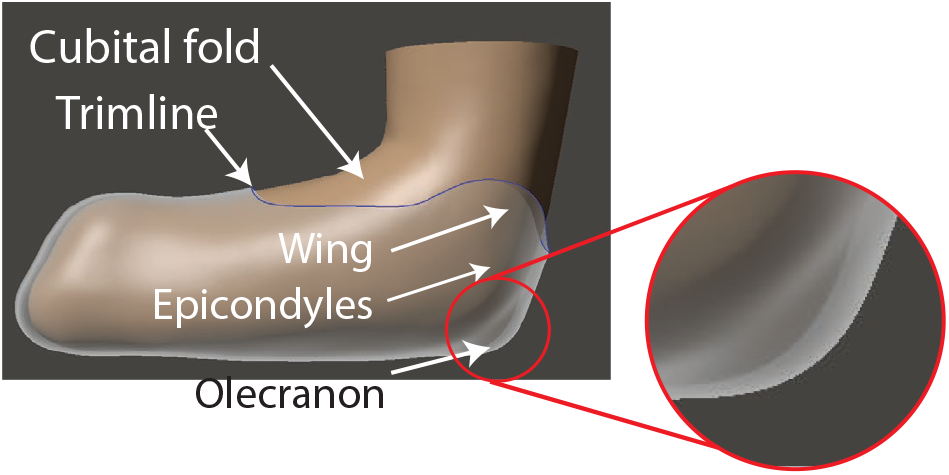
A diagram showing key areas of upper-limb anatomy and socket terminology. The olecranon is highlighted for clarity.

Steps five, six and seven are optional, but can lead to a better fitting socket. Sometimes, when check sockets have been used as detailed in step five, prosthetists can combine steps eight and nine when fabricating the socket.

### 2.4 Digital build of sockets

We attempted to replicate steps 1-5 of the example *traditional* procedure using digital tools. All experimental procedures and decisions regarding design and manufacturing were carried out by the engineering team (Co-authors: Olsen, Dupan, Nazarpour and Dyson), who did not have any formal training in prosthetics. Expert advice was sought from a professional prosthetist (Co-author: Day) to minimise the likelihood of harm to the participants.

We utilised a low-cost optical scanner (*circa*. £300, Sense v1, 3D Systems, USA) to capture the shape of the limb. Optical scanning is a non-contact process hence limb socks were not used. The scanner and software combination supported geometry-only scans, so limb markings were not required.

The Strathclyde Supra Olecranon Socket (SSOS), was chosen to be made for all participants. The method was developed to suit a wide variety of trans-radial amputees. In this method, suspension is achieved by gripping above the olecranon. The wings that enclose the epicondyles are mainly for stability and to provide secondary suspension. For a SSOS socket, the residual limb should be cast with the elbow flexed at an angle of 90° to ensure that adequate socket suspension is achieved above the olecranon.

Participants were instructed to hold their limb still at a right angle and to not rotate their arm from a natural resting position. Three participants required reflective scanning markers adhered to their limb to obtain complete scans, e.g. Fig. 2(a). Markers assist the scanner by providing points of reference, especially useful for limbs without distinct texture or features. Up to four attempts were made until a scan that appeared free from major flaws was obtained. The scan with the least flaws upon visual inspection was selected for further processing.

**Figure 2:**
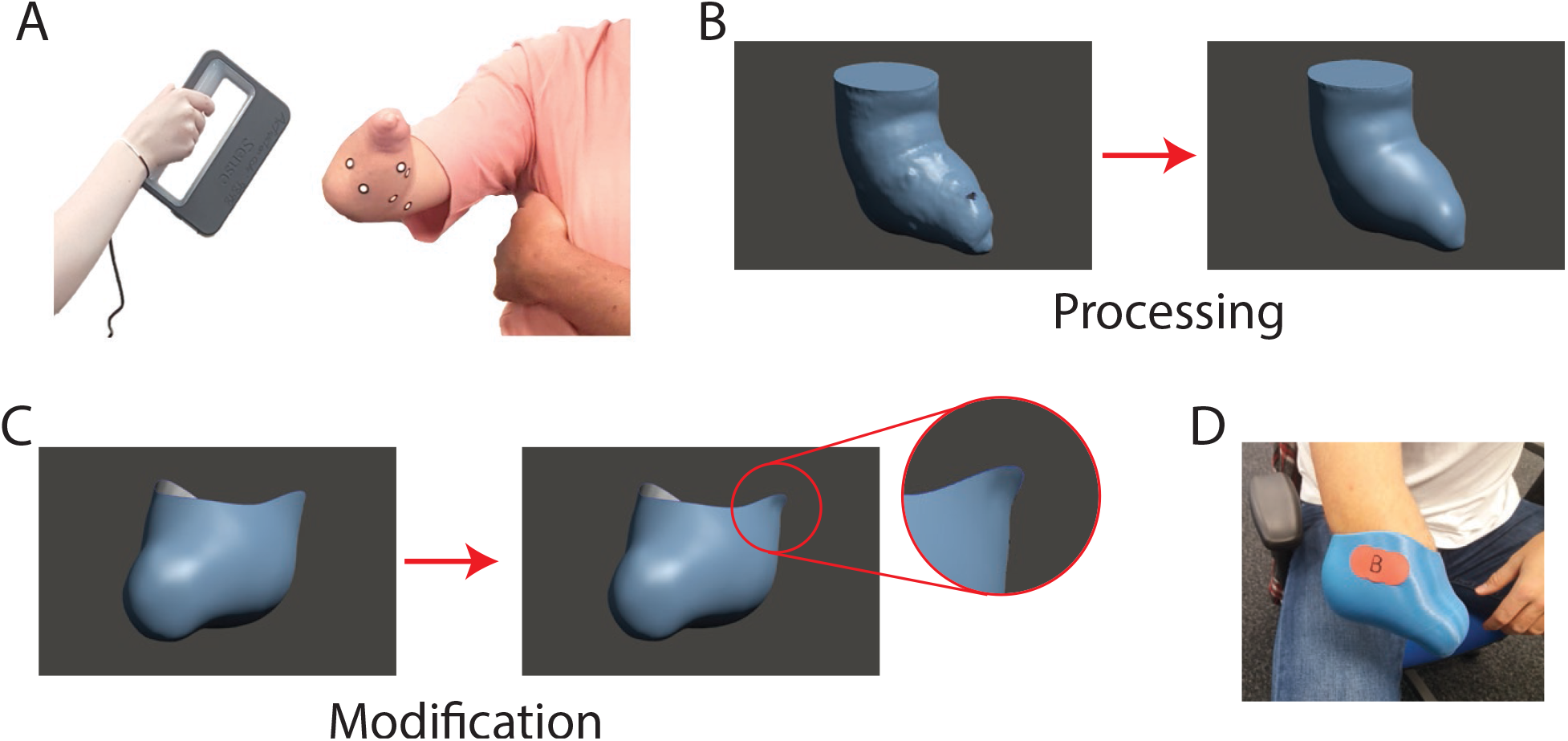
A: A participant with reflective scanning markers on their limb; B: A scan of a participant’s limb before and and after processing; C: An example of a unmodified and modified socket with contouring above the epicondyles; F: The same participant wearing an unmodified socket.

Two socket types were made for each participant: 1) unmodified, made directly from the scan with no contouring; and 2) modified, altered in CAD (Computer-Aided Design) software by the engineering team to approximate the modifications a prosthetist would make, based on the suggestions of the clinician. Specifically, these modifications simulated the application of the moulding grip and the post-casting sculpting of the postive cast. For participants with moderate-severe pain, phantom sensations or skin sensitivities several pairs of sockets were built with subtle differences in contouring and geometry to reduce the risk of the participant experiencing discomfort.

The scans were processed in Autodesk Meshmixer to remove artifacts, patch holes and perform smoothing. A comparison between a pre-processed and post-processed scan is shown in Fig. 2(b). The scans were visually inspected to locate the approximate area of reference points such as the epicondyles, cubital fold and olecranon. These areas were used to guide the trimlines, which were drawn by hand in Meshmixer. These sockets were saved as the unmodified design. The unmodified socket from each participant was used as the base file for the ‘modified’ socket to keep the geometry consistent. If multiple pairs of sockets were required, different trimlines were drawn and saved as separate files. Between each participant’s different pairs of sockets the only difference was the trimline and wing height, and within each pair the only difference was whether it was modified or unmodified. Fig. 2(c) shows an example for the difference between an unmodified and modified socket. The contouring above the epicondyles is highlighted.

The 3D models were converted to meshes in Autodesk Recap and thickened to 4mm in Autodesk Inventor. Ultimaker Cura and Ideamaker were used to slice the models prior to printing. The sockets were printed on three types of FDM printers: Ultimaker 2+, Ultimaker 3 and the Raise 3D Pro 2 Plus. The materials used were generic PLA (Polylactic Acid), PLActive (antibacterial PLA/Copper Nanocomposite) and Taulman Gu!deline medical grade PET-G (Glycol Modified Polyethylene Terephthalate). The sockets were printed using a 0.4mm nozzle at a layer height of 0.18mm, infill 15% at a 45 degree angle relative to the print bed to minimise support material. The print speed and temperature varied with the filament, as per the manufacturer instructions. The printer model, material used, software and printer settings were kept consistent between each pair of modified/unmodified sockets to allow a fair comparison. Some pairs of sockets required light manual sanding due to *stringing* during printing.

### 2.5 Interview

All participants tried their sockets. An example is shown in Fig. 2(d). Participants were presented with both modified and unmodified sockets, marked A and B, respectively. Participants were not aware which had been modified. The order in which the sockets were tried was randomised. Participants with multiple pairs were asked to try on all A or all B sockets, depending on the testing order for their session, and pick the socket type that fitted them best. Other sockets were discarded. All participants took part in an interview. The interview included a set of questions but the discussion was kept open-ended. The questions were:

1. How comfortable do you find the socket? What are your initial thoughts?
2. How secure do you find the socket? Are you concerned about the socket slipping off your arm?
3. Are there any areas of the socket you would like to change? How do you feel about the trimlines? The wings?
4. How long do you think you could wear the socket without discomfort?
5. How do you feel about the socket compared to any of the sockets you currently use (if any)?

Following the interview, participants were informed which socket was modified and which was unmodified. Participants were then invited to try on the sockets again, with and without limb socks. Some participants chose to try their own sockets on for comparison.

## 3 Results

In this section, we first report on the cost and time investment required to manufacture the sockets with digital methods. Then, we provide a summary of the feedback from participants.

### 3.1 Quantitative analysis

Scanning took approximately 2-15 minutes per limb, depending on the participant and lighting conditions. CAD modifications, from raw scan to finished pair of modified/unmodified sockets took approximately 30 minutes. All six participant scans resulted in 3D printed diagnostic sockets. Print times varied between 12hr for the smallest socket to 17hr for the largest using the print settings for PLA, with an average between participants of appproximately 15hr. Post-production removal of support material and manual sanding took approximately 5 minutes per socket. Including the discarded support material, the mean socket weight between participants was 93.6g. The material cost of each socket was *≈*£3 when using PLA filament, *≈*£9 for PLActive, and *≈*£10 for Guidel!ne.

### 3.2 Summary of the interview responses

Of the six participants, four participants P1-4 completed the feedback session. In the following we summarise the comments. The texts in the *italic* font are direct quotes from the participants. Clarifications were sought from the participants during interview for ambiguous answers. The participants’ comments were used to create categorical data. Key results are summarised in Fig. 3. The raw interview notes are accessible via the supporting material.

**Figure 3:**
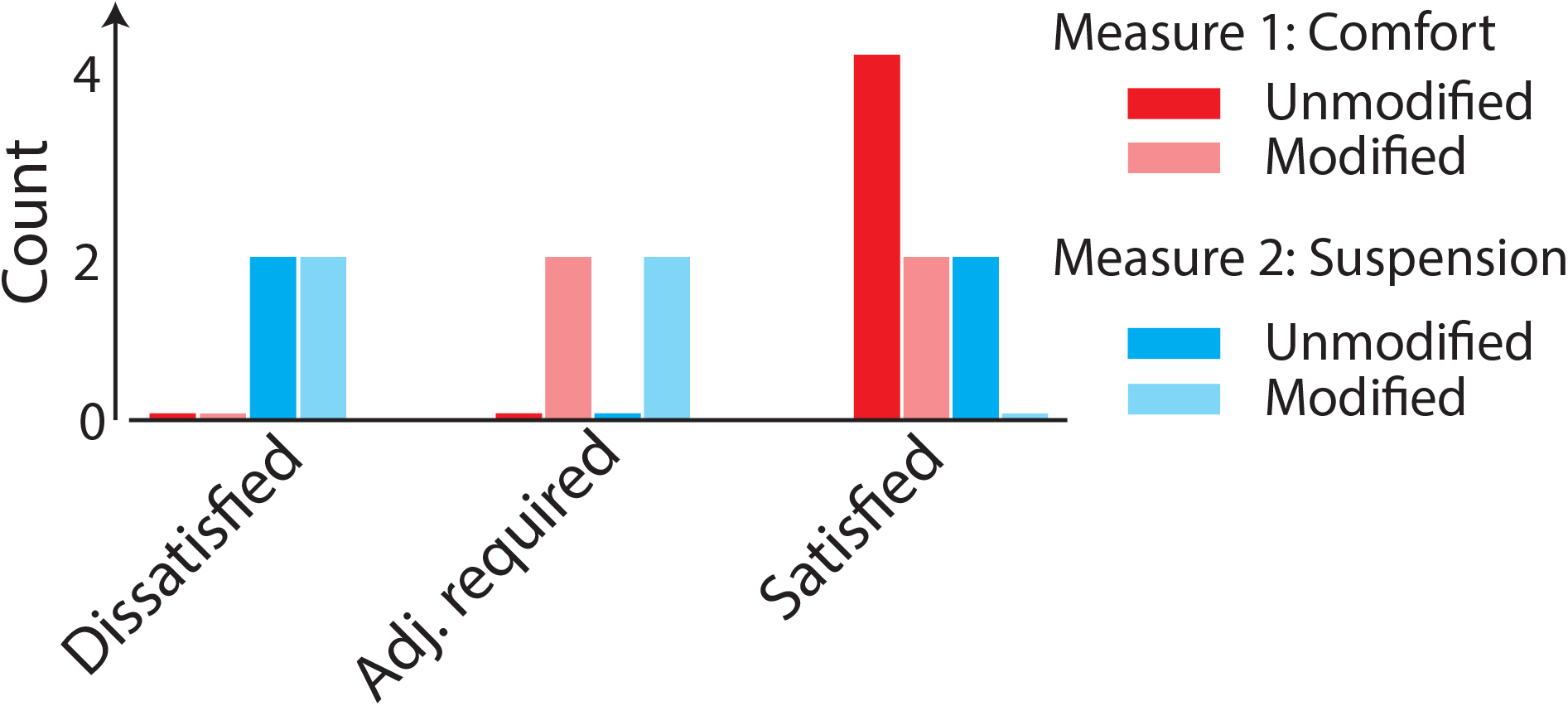
A summary of how satisfied the participants were with the key factors affecting socket fit: comfort and suspension.

#### 3.2.1 Comfort

All four participants found their unmodified socket *very comfortable*. The feedback from the modified sockets was slightly less favourable, with only two participants finding their modifed socket very comfortable. The other two participants highlighted areas which could be improved. Participant P1 stated that their unmodified socket was:

> … the most comfortable socket I’ve ever tried.

Overall, comfort was rated extremely well from the feedback, with only minor problems being highlighted.

#### 3.2.2 Suspension

The suspension of the sockets was generally unsatisfactory, with only two participants finding one of their sockets (both unmodified) to be secure. For instance, participant P1 achieved excellent suspension, and felt that they could:

> … suspend my entire bodyweight with this (socket).

Other participants were not satisfied by the achieved suspension. For example discussing both their modified and unmodified sockets, participant P3 stated that:

> It’s not secure. I can take it off easily.

Despite neither type of socket achieving the majority of participants feeling secure, most participants did feel more secure in their unmodified socket than the modified version.

#### 3.2.3 Socket modification problems

All participants found their socket comfortable. Nevertheless, participants P3 and P4 both mentioned the trimline above the olecranon of both of their sockets could be lower, with participant P4 stating:

> … it’s not uncomfortable, just different to what I usually use.

Similarly, both participant 1 and 2 felt that the inner wing of their modified sockets was tight, with participant 2 stating:

> … the inside wing causes a bit of friction, which if I wore it for a long time might get sore.

Overall, the localised problems highlighted were all minor and related to the wings, trimlines and proximal contouring.

#### 3.2.4 Tolerability

When asked how long they believed they could wear their sockets for without causing discomfort, participant P3 stated *all day*, for both their unmodified and modified socket. Participant P1 and P4 stated they could wear their unmodified socket all day, but were unsure about their modified socket. Similarly, participant P2 was unsure about either of their sockets, due to the inner wing.

#### 3.2.5 Comparison to own socket

When comparing the 3D printed sockets to their own, participant P2 and P3 preferred their usual socket to both of their 3D printed sockets. Participant P1 preferred their unmodified 3D printed socket to any they had ever owned, and participant P4 was unsure how to compare the sockets as they were different to what they were accustomed to.

#### 3.2.6 Additional comments

Some participants chose to share further comments after the structured interview session had concluded. Participant P1 stated that:

> … I prefer this (method)… some of my sockets have required so many refitting sessions and visits to the clinic… it took so much time… you have demonstrated that you can create a socket that fits perfect and has all the requirements including load bearing and has very little or no pressure points.

Several participants expressed they were impressed at the level of detail achieved using the low-cost digital scanner when shown their scans on screen.

#### 3.2.7 Results summary

Overall, only participants P1 and P4 found at least one of their sockets to be both sufficiently comfortable and secure. Participants P2 and P3, despite generally finding their sockets comfortable, did not find the suspension to be adequate, and suggested minor changes which they thought would improve the fit if another iteration of sockets were to be trialled. This is summarised in Fig. 4.

**Figure 4:**
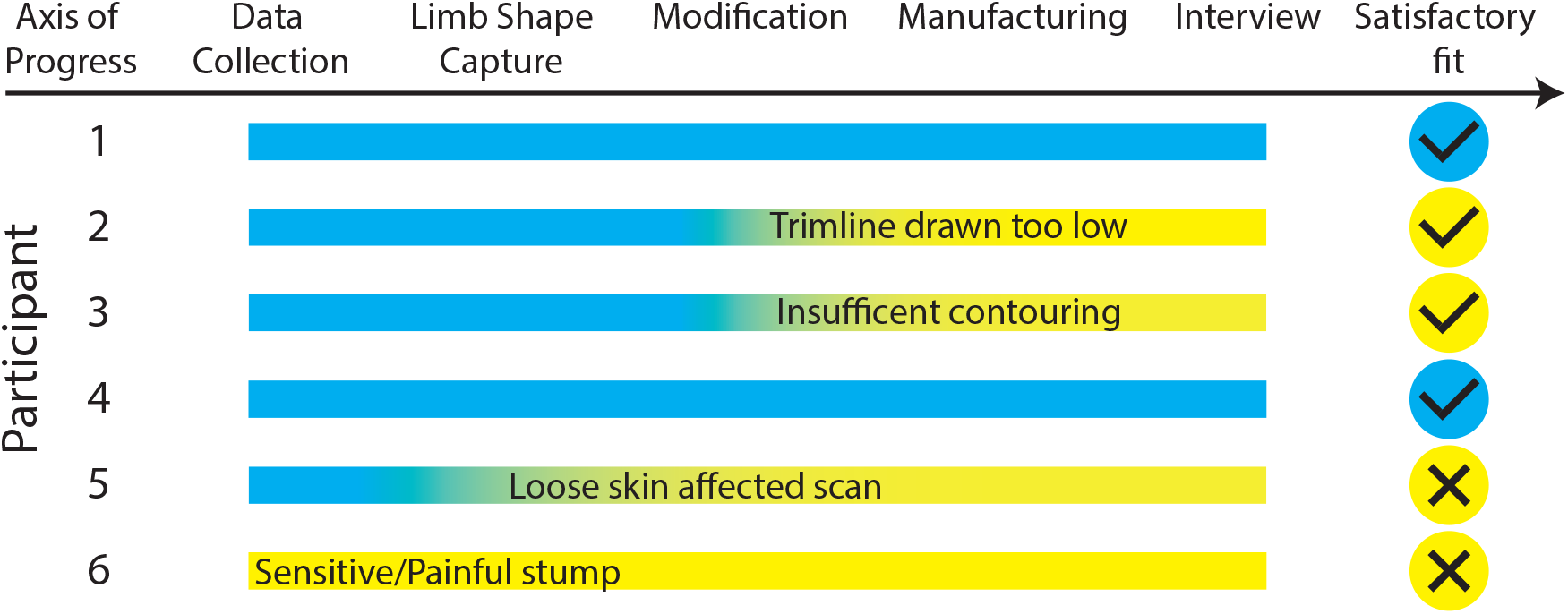
A timeline highlighting the various stages of manufacturing that negatively impacted the fit of the sockets and the overall satisfaction levels of the participants.

## 4 Clinician commentary

Lack of clinical experience within the engineering team appeared to contribute significantly to the cases where participants were not satisfied with their sockets. Participants P2 and P3 both cited the wings, trimlines and proximal contouring to be the main problem. As with the traditional workflow, very small changes to the digital modification process can produce significantly different outcomes regarding comfort and security. Modifications are usually based on experience, and often the *feel* of the limb is an important factor when determining how to modify the socket. These types of methods are particularly difficult to translate to a digital workflow, especially with novice operators. Participant P5’s sockets failed because of loose skin on the residual limb. It is probable that this could have been remedied using a limb sock. However, it remains the case that optical scanning alone can be vulnerable to failure when capturing nuanced physical features which are necessary for good fit. In the case of participant P6, a particularly sensitive area around their epicondyles was not reported during the interview and therefore was not accounted for during the diagnostic socket modification. This would have required the socket to be modified or re-made regardless of whether the method was digital or traditional.

## 5 Discussion

In this section we discuss the learning from this study in the wider context of modern methods for socket manufacturing. Finally, we explore the limitations of the study and future avenues for research.

### 5.1 Analysis of the experimental results and the interview

We provided a realistic, digital adaptation of the traditional method to manufacture upper-limb sockets. Low-cost systems were utilised because using high barrier-to-entry technology or excessive software sequences would negate two of the key benefits of switching to modern technology, namely simplicity and reduced costs. The sockets were tested by people with limb difference and feedback was collected regarding comfort and suspension. Expert clinical opinion was sought to discuss the outcome of the study and highlight why problems occurred. Our results confirmed that modern technology is vulnerable to the same problems as traditional socket manufacturing when used without expert operation. This supports anecdotal evidence that adopting a naive plug-and-play approach using optical scanning and 3D printing for prosthetics is unlikely to produce satisfactory results [13].

Often, the main argument presented for embracing 3D printing within prosthetics is that it is faster, cheaper and produces better results than traditional methods [7, 47]. Due to the lack of documentation of practices in standard prosthetics and orthotics clinics, quantifying any time or cost savings using current digital methods is difficult. The approximate timeline of our digital socket manufacturing method compared to the traditional is shown highlighted in blue in Fig. 5. Assuming no errors occur, the methods require similar timescales - the key benefit offered is reduced reliance on manual labour, particularly in the case of re-fits requiring new plaster casts. It is reasonable to assume that 3D print speeds will increase, as such digital workflows still offer the potential for same-day prostheses, which would be a huge breakthrough for the prosthetics industry.

**Figure 5:**
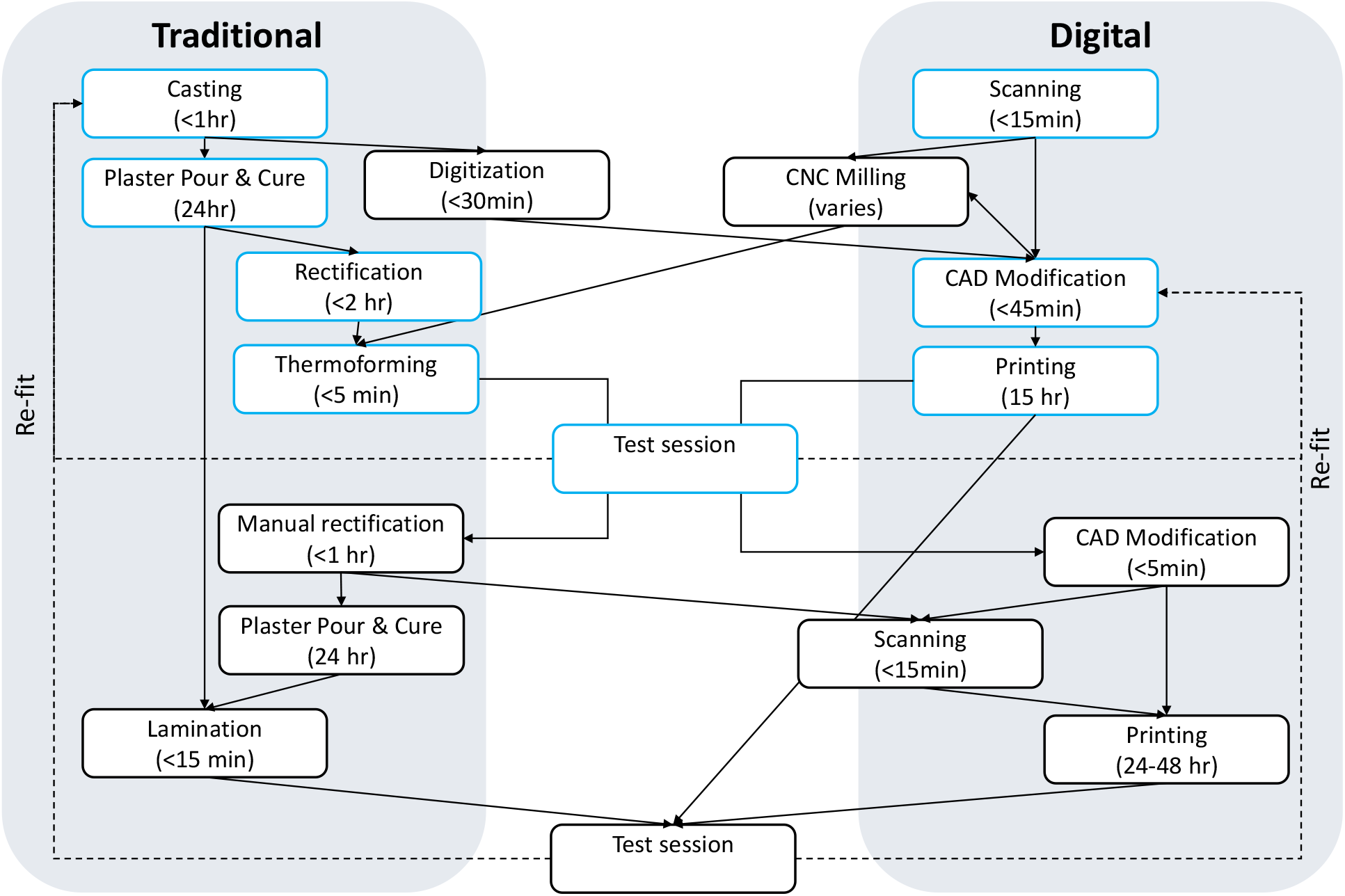
Approximate timescales of average traditional and digital socket manufacturing workflows. The steps compared in the practical element of this paper are highlighted in blue.

Usually, any suspension or contouring problems would be rectified whilst testing the diagnostic socket with the patient in clinic. Besides trial and error, there would be only two ways to carry out this rectification using digital tools. First, the 3D printed diagnostic socket would need to be heated and re-contoured as per the usual procedure. This is possible as PLA can be heated and reformed in the same manner as thermoformed sockets. Then, to replace the second plaster pour either: 1) the same modifications made to the socket would be approximated on the CAD model or 2) the inside of the modified diagnostic socket scanned and digitised, as shown in Fig. 5. Regardless, a clinician’s expertise is crucial during this step, and arguably it would be more efficient using the traditional method. This further demonstrates the need for expert clinical intervention, irrespective of the method used.

### 5.2 Further work and limitations

The scanner and software used did not offer the option to view the colour of the scans. Hence, creating the trimlines and wing shape was particularly difficult. Marking the key areas of the limb is an essential step in guiding the trimline cut. Use of alternative technology would have allowed the colour from the scans to be utilised. A future study with a scanner and software combination that captures colour operated by an experienced prosthetist would allow for closer comparison to the current standard.

Despite the majority of participants finding their sockets very comfortable, only two participants also achieved socket security. Indeed, achieving socket comfort without suspension is essentially useless, and vice versa. Future research should be conducted to investigate how improved suspension within upper-limb sockets can be achieved using digital methods.

No terminal device was attached whilst the participants were wearing the diagnostic sockets and they were only worn for a short duration of time, up to 20 minutes. This meant that the comfort and security whilst bearing weight was not tested. A future comparison will assess the socket when used with a terminal device, bearing load at a variety of angles. Additionally, the safety of the sockets when bearing load would need to be considered, potentially using stress modelling or mechanical testing. Only the SSOS socket was adapted into the digital workflow for this study. Future studies should consider other styles which may be better suited to translation to a digital workflow. The length of the residual limb is an important factor in deciding which socket style to use, and contributes to the overall success of the socket.

## 6 Outlook

In the following section we will discuss the contrasting perspectives of the research and clinical communities, the safety and legality of 3D printing and the potential benefits moving forwards.

### 6.1 Research perspective

There is a lack of documentation detailing the intricate procedures carried out in clinics, and the way that clinicians make prostheses is rarely studied [14]. Often techniques are based on personal experience, views and implicit knowledge [11, 16, 30, 36, 40], which leads to inconsistencies in methods [36]. Additionally, prosthetists have protected profession status in 21 countries in Europe, with similar regulations globally [2,10, 42]. Protected profession status means that only registered individuals can create and distribute functional prostheses in regulated areas [2, 10, 26, 42]. Although important for safety, this status likely contributes to a lack of accessible information for researchers developing technology intended to assist clinicians. Together, these factors make accessing and understanding highly skilled procedures difficult for researchers outside of the clinical domain.

### 6.2 Clinical perspective

Prosthetists see significantly fewer people with upper-limb absence relative to lower-limb [1, 9], and each has unique characteristics and needs. It is unsurprising that current ISPO regulations do not define one specific method must be used use in order to create devices [16]. Hence, procedures are difficult to capture in a digital workflow. Clinicians looking to adopt digital techniques within their practice have two options: devise their own workflow or buy in to a pre-made P&O software package. Conventional CAD software suites are not designed specifically to manufacture prosthetics, as such they are highly flexible but also complex. Alternatively, companies offer proprietary software targetting P&O clinics, such as the CanFit™ software developed by Vorum [44], or WillowWood’s OMEGA™ [46]. Despite offering a simplified solution, these programs often have a high-barrier to entry in terms of cost, with limited published research investigating their capabilities and limitations when used for upper-limb populations.

Further research is also needed to clarify how the properties of additive manufactured sockets behaves. Currently, nylon-resin sockets are reliable and predictable in terms of stiffness and other properties. Clinically validated mechanical testing of a variety of materials and designs are required to ensure prosthetists can use digital methods with confidence.

### 6.3 Safety and legality concerns

Limited evidence is available regarding the durability of prosthetic sockets [6] and multiple factors affect the mechanical properties of the final device [3, 6, 18]. Catastrophic failures of 3D printed lower-limb sockets have been documented [35]. Standardised safety and durability tests exist for lower-limb devices [17], which has enabled promising results from 3D printed trans-tibial sockets [6, 29]. In contrast, no such standard exists for upper-limb sockets [23]. This makes introducing new materials difficult, as it is difficult to draw comparisons against traditional materials. It is not clear whether digital scanning is as accurate as traditional techniques [22] and the relationship between socket fit and comfort remains unclear [39]. Additionally, digital methods are not required to be taught at P&O centres under ISPO guidance [16] posing challenges for translating research into clinical practice.

### 6.4 Potential Benefits

Digitisation offers numerous potential opportunities outside of the manufacturing process. Unlike traditional methods, digitised scans and designs can be easily stored and shared near instantaneously via the internet. These benefits would allow clinicians and users to obtain input from colleagues and or specialists, enabling a more uniform quality of care irrespective of location. When enhanced with documentation detailing the outcomes of individual prosthetic interventions, shared digital data could have significant value. Such data would enable prosthetists facing an unusual or challenging case to search for precedents, linking them with relevant cases and colleagues and enabling solutions to be found more efficiently. Digital captures of prosthetics interventions over time would provide quantifiable data, which could potentially train algorithmic approaches in the future. A data format also enables use of algorithmic tools such as finite element analysis, which has been proposed to improve loading conditions within sockets [24].

## 7 Conclusions

In this study, a fully digitised, low-cost method of producing transradial diagnostic sockets was presented. Volunteer feedback and expert commentary was sought. The contrasting perspectives of the research and clinical communities were discussed, alongside safety concerns and the potential benefits of adopting digital methods for socket creation. The outcomes of the study confirmed that clinical expertise is crucial for creating well-fitting prosthetic sockets, regardless of the methods used. The discussion highlighted the need for further safety testing of 3D printed prostheses to help clinicians make informed decisions as to how and whether to adopt digital manufacturing into their clinics. Further collaboration is required between the clinical and research communities to form digital tools that are useful for patients and clinicians.

## Supporting information

Supplementary_data_Participant_Info

Supplementary_data_Qualitative

Supplementary_data_Quantative

## 8 Acknowledgements

The authors would like to thank Dr. John Head of Salford University for his helpful comments on an earlier version of this paper.

## Notes

### Competing Interest Statement

The authors have declared no competing interest.

