## Supplementary_data_Participant_Info for "3D-Printing and upper-limb prosthetic sockets; promises and pitfalls"

| Participant | Age Bracket | Gender | Amputation | Prosthesis Use | Notes |
|---|---|---|---|---|---|
| 1 | 50-60 | M | Acquired - transradial | Uses several sockets for different applications with static (passive adjustable) attachments - i.e. sports, driving. Doesn't usually wear a prosthesis for leaving the house. Previous experience using a myoelectric device. | Phantom sensations at distal amputation site and minimal pain. Pain is worse in the cold. |
| 2 | 40-50 | F | Acquired - transradial | Uses several sockets for different applications with static and body powered attachments - i.e. cooking, driving. Doesn't usually wear a prosthesis for leaving the house. Previous experience using a myoelectric device. | Occasional phantom sensations and pain at distal amputation site. |
| 3 | 30-40 | M | Acquired - transradial | Myoelectric prosthesis used daily for their job. | No pain or sensations reported. |
| 4 | 20-30 | M | Congenital - transradial | Various sockets (body powered attachments) for specific tasks such as going to the gym, driving. Doesn't usually wear a prosthesis for leaving the house. | N/A as congenital. |
| 7 | 60+ | M | Acquired - transradial | Various sockets (body powered/static attachments) used for specific tasks. | No pain or sensations reported. |
| 6 | 60+ | M | Acquired - transradial | Various sockets with body powered and static (passive adjustable) attachments for specific tasks such as craft activities. Usually wears a passive anthropomorphic prostheses when leaving home. Previous experience using a myoelectric device | Severe nerve damage at distal end of limb. Constant pain and phantom sensations. |
