## Supplementary_data_Qualitative for "3D-Printing and upper-limb prosthetic sockets; promises and pitfalls"

| | | Q1: How comfortable do you find the socket? What are your initial thoughts? | | Q2: How secure do you find the socket? Are you concerned about the socket slipping off your arm? | | Q3: Are there any areas of the socket you would like to change? How do you feel about the trimlines? The wings? | | Q4: How long do you think you could wear the socket without discomfort? | | Q5: How do you feel about the socket compared to any of the sockets you currently use (if any)? | | Q6: Additional quotes |
|---|---|---|---|---|---|---|---|---|---|---|---|---|
| **Participant** | **Order** | **UM** | **M** | **UM** | **M** | **UM** | **M** | **UM** | **M** | **UM** | **M** | **** |
| 1 | UM First | Very comfortable, easy to don. I think it's actually the most comfortable socket I've ever tried. | Comfortable but slightly harder to don than UM due to inner wing. | This feels very secure, I could suspend my entire bodyweight with this (referring to the socket) | This one doesn't feel as secure as the first* (*first = UM) | None. | The inner wing causes some pressure so I'd probably change that. | All day, it's very comfortable. | I'm unsure because of that inner wing. | It's extremely comfortable and I'm really happy with fit. It's the best socket fit I've ever tried on. | This one is not as good as the first because of the wing. | Personally, I prefer this (referring to the method) for getting a socket. Some of my sockets have taken so many refitting sessions and visits to the clinic, when you add it up it took so much time. I am totally amazed with the results of these sockets. As I showed you the prosthetic I have for weight lifting prosthetic and is not ideal and that was after 3 visits with time and cost and I think what you have demonstrated is that you can create a socket that fits perfect and has all the requirements including load bearing and has very little or no pressure points. (Participant also stated they were impressed with the quality of the scan when shown their scan on screen) |
| 2 | UM First | Very comfortable, the extension range is good, the length is good. It's a bit snug at the end* so pressure might be a problem long term but it feels comfortable. (*end = distal amputation site) | This feels the same as the first one* except it feels looser. (*first = UM) | It's not secure, I can remove it quite easily (referring to the socket), it's a bit loose around the elbow because the trimlines are low, they need to be about 1cm higher, it's not far off though, maybe not even 1cm. | This one is looser than the first, it actually feels lower too. | As I said, the trimline could be higher but it's not far off, 1cm or less higher on the wings would be ideal. | With this one, the inside wing causes a bit of friction, which if I wore it for a long time might get sore. | I'm not sure. | It's the same as the first socket, the wing might cause problems if I wore it for a long period of time. | I prefer my usual socket as I can rotate in it really well, I use it for driving so I find that useful. I'd usually wear socks under my socket too. | Same as the first, except this one is looser so the first socket was better out of the two. | Both the sockets were better with a sock, except then they felt a bit snug. Overall they really weren't far off being a good fit, I think with small changes the fit would be quite good. (Participant also stated they were impressed with the quality of the scan when shown their scan on screen and also commented on how many detailed the sockets were in comparison to others they had received.) |
| 3 | M First | Same as the first*, feels very comfortable, no issues. (*first = M) | It feels very comfortable, no issues with it. | This isn't secure either, but the first was less secure. This one is a bit better in terms of grip. I can still take it off quite easily though. | It's not secure, I can take it off easily. | Same as the first really. | The left wing needs a bit of modification and the back is higher than my usual (socket) but not bad. I do like the socket and I find it extremely comfortable but I'd be worried about the security, I think it's because of the low trimline. | All day. | All day. | Same as the first. | I'd say it's similar to my own socket but slightly less preferable due to security issues. | (Participant stated they were impressed with the quality of the scan when shown their scan on screen, particularly the details that could be captured.) |
| 4 | M First | Same, very comfortable. | Very comfortable | This one feels secure. | It doesn't feel completely secure. | Same (as the first*). (*first = M) | Not really, the trimline at back could be lowered but it's not uncomfortable, just different to what I usually use. | All day. | I'm not sure. | Same as the first. | Not sure because it's different to my usual, not bad, just different, so it's hard to compare. | None. |
| 5 | M First | Sockets were much too loose to gauge comfort or security (i.e. not in full contact with the limb) so the feedback session was terminated. | | | | | | | | | | |
| 6 | UM First | Feedback session terminated as participant divulged that a channel is usually created to relieve pressure over their inner epicondlye (due to nerve damage) after the sockets were manufactured. | | | | | | | | | | |
