## Supplementary_data_Quantative for "3D-Printing and upper-limb prosthetic sockets; promises and pitfalls"

| Participant | Print time (PLA Settings) | Socket weight incl. support material (grams) | Cost (£) - PLA | Cost (£) - PLActive | Cost (£) - Gu!deline |
|---|---|---|---|---|---|
| P1 | 16:18:00 | 103.30 | 3.35 | 10.25 | 10.93 |
| P2 | 14:27:00 | 87.70 | 2.84 | 8.70 | 9.28 |
| P3 | 14:16:00 | 92.00 | 2.98 | 9.13 | 9.74 |
| P4 | 12:01:00 | 78.00 | 2.53 | 7.74 | 8.25 |
| P5 | 17:36:00 | 107.40 | 3.48 | 10.65 | 11.37 |
| P6 | 14:59:00 | 93.10 | 3.02 | 9.24 | 9.85 |
| Mean | 14:56:10 | 93.58 | 3.03 | 9.28 | 9.90 |
